# Ageing, TOR and amino acid restriction: a cross-tissue transcriptional network connects GATA factors to *Drosophila* longevity

**DOI:** 10.1101/036848

**Authors:** Adam J Dobson, Xiaoli He, Eric Blanc, Ekin Bolukbasi, Yodit Feseha, Mingyao Yang, Matthew DW Piper

## Abstract

Animal lifespan can be extended by dietary restriction (DR), but at a cost to fitness. This phenomenon depends on essential amino acids (EAAs) and TOR signalling, but roles of specific tissues and downstream transcriptional regulators are poorly characterised. Manipulating relevant transcription factors (TFs) specifically in lifespan-limiting tissues may ameliorate ageing without costs of DR. Here we identify TFs which regulate the DR phenotype in *Drosophila*, analysing organs as an interacting system and reducing its transcriptional complexity by two orders of magnitude. Evolutionarily conserved GATA TFs are predicted to regulate the overlapping effects of DR and TOR on organs, and genetic analyses confirmed that these TFs interact with diet to determine lifespan. Importantly, *Srp* knockdown insulated fly lifespan from the pernicious effects of EAAs, but tissue-specific knockdown reduced the corrolary costs. These results provide the first indication that benefits of EAAs for early-life fitness can be decoupled from longevity by tissue-specific transcriptional reprogramming.

## Background

How can lifelong health be maximised? Answering this question is a major goal, as ever-increasing human lifespans outpace advances in gerontology, at great social, personal and financial cost^1^. Dietary restriction (DR) has the evolutionarily conserved capacity to improve lifelong health by reducing nutrient intake, but at a cost of reduced biological fitness and vigour in youth^2^. Despite having been discovered 80 years ago^3^, the molecular mechanisms underpinning lifespan extension by DR remain elusive. Defining these mechanisms could ameliorate the burden of ageing without the costs of DR.

Calories do not fully account for the benefits of DR: specific nutrients and their relative ratios are key^4–6^. In *Drosophila*, the ratio of dietary sugar to yeast modulates lifespan, which is explained by essential amino acids (EAAs) from the yeast^7^, and importantly the same mechanism is conserved in mice^8,9^. Recent evidence indicates that the phenotype of EAA-restricted *Drosophila* is recapitulated by pharmacologically suppressing the Target of Rapamycin (TOR) pathway^10,11^, consistent with molecular evidence that EAAs positively regulate TOR^12^. Understanding of how TOR curtails lifespan is incomplete, although maintenance of proteome quality likely plays a role^13–15^. TOR also affects transcription^16–18^, but to date this effect has been relatively poorly studied.

In *Drosophila*, transcriptomic responses to DR and TOR have been characterised at the cellular and organismal levels^18^, but information on tissue-specific and organ-specific transcription is a requisite advance for many reasons. Primarily, an animal’s overall phenotype is determined by tissues coordinating to match their collective functions to the environment. Consequently, to fully understand organismal phenotypes we must account for tissues as an interacting and integrated system. By quantifying the changes to that system in low-TOR or DR states, we gain a complete view of how tissues collectively mediate the DR phenotype. Such an understanding is likely to help identify manipulations of regulators of the DR transcriptional state, which may impart longevity even under *ad libitum* feeding. Minimising the number of tissues in which these regulators are manipulated may ameliorate associated costs, if costs and benefits of DR are mediated by distinct tissues. This approach recognises the findings of studies of insulin signalling, which indicate that tissue-restricted manipulations are sufficient to extend lifespan, and therefore that transcriptional mechanisms affecting longevity are incompletely described by organismal analyses^19,20^. Altogether, these observations suggest that understanding longevity via DR requires (1) modelling of transcription across systems of interacting tissues, (2) identification of changes to that system under DR, (3) quantification of the TOR-dependence of these changes, (4) prediction and testing of tissue-specific regulators of DR-like gene expression.

Recent analytical advances facilitate prediction of transcriptional regulators from patterns of gene expression. Regulatory elements, such as transcription factors, can be predicted by enrichment analysis amongst groups of genes^21,22^. Transcriptional network analysis, also known as coexpression analysis, identifies transcripts which are expressed together^23,24^. The patterns in a transcriptional network therefore reflect the tendency of genes to be switched on and off together, which can be considered a reflection of the underlying gene-regulatory machinery. Consequently, transcriptional network analyses provide a strong basis from which to predict regulatory elements. Network analysis can also answer the need to understand tissues as a complete system, by identifying how patterns of expression in one tissue which correspond to that in others. Therefore, by defining transcriptional networks across tissues and how they change under DR, we can distill the complex interactions between transcripts and tissues that define the phenotype of the DR animal, and systematically reflect the status of the underlying gene-regulatory machinery.

Here, we address a key question: By studying transcription across tissues, can we identify manipulations of transcriptional regulators to recapitulate the DR phenotype? This study is facilitated by dietary manipulations that offer a precise tool-set to dissect DR. Previously, we developed a semi-defined *Drosophila* diet, which is optimal for early-life egg laying, in which 50% of available EAAs are provided as a supplement to yeast-based medium^7,10^. Against this “fully fed” control, lifespan can be extended by two interventions, both with correlated fecundity costs to egg laying in early life: either omission of the EAA supplement (i.e. DR); or the addition of rapamycin (EAA+rapamycin), which extends lifespan in the presence of EAAs by suppressing TOR pharmacologically. These tools allow us to establish the effects of EAA dilution, and the TOR-dependence of these effects. Taking these diets and capitalising on orthology between *Drosophila* and vertebrate organs, we studied transcriptomes in the brain, fat body (the analogue of the vertebrate liver and adipose), gut, ovary and thorax (which largely comprises muscle). We then used transcriptional network analysis to analyse these tissues integratively, reflecting underlying regulatory machinery. Promoter analysis was then applied to identify candidate transcription factor regulators. Crucially, when we tested these transcription factors genetically, we were able to validate their role as mediators of dietary effects on lifespan, and also demonstrate that costs of lifespan extension can be mitigated by targeting interventions to specific tissues.

## Results

The effects of DR on ageing are thought to be mediated by the TOR pathway. Here, by characterising the organ-specific transcriptional changes caused by EAA restriction (DR) and pharmacological TOR suppression, we isolate changes that are associated with longevity in both experimental conditions, and address whether the effects of DR mimic the effects of low TOR. We also associate *cis-*regulatory elements with these changes, to predict relevant transcription factors. The study design is presented in Figure 1. We term DR as a diet-induced longevity condition, and rapamycin supplementation as a drug-induced longevity condition.

**Figure 1.**
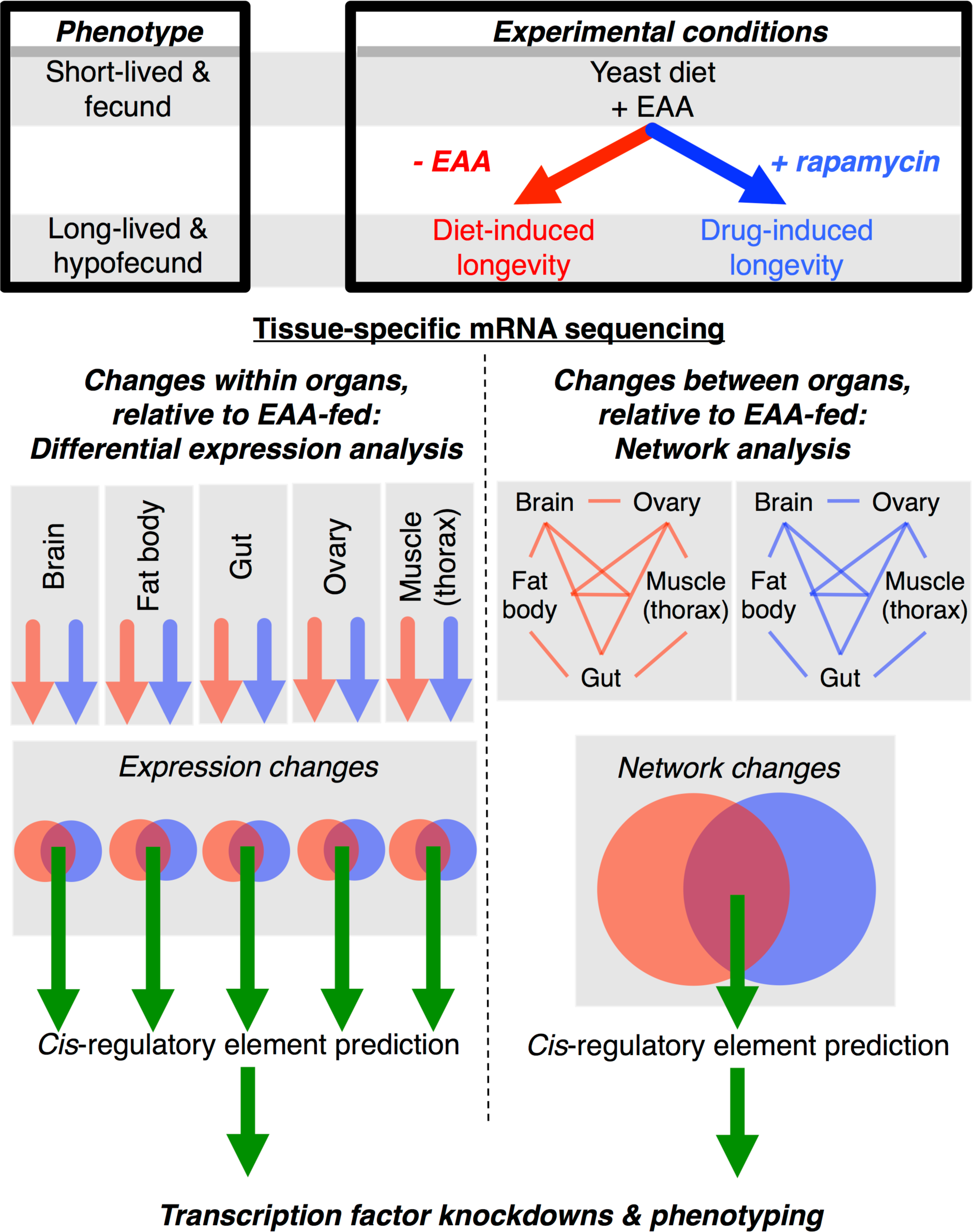
Study design. Against a control in which short lifespan is determined by enrichment of essential amino acids (EAAs), lifespan is extended by DR of the EAAs (diet-induced longevity), or pharmacologically, by administration of the TOR-suppressive drug rapamycin (drug-induced longevity). This lifespan extension comes at a biological cost of reduced early-life fitness (i.e. fecundity). Tissue-specific transcriptional changes associated with both conditions are therefore longevity-associated, EAA-dependent and TOR-dependent. Identifying *Cis*-regulatory elements associated with these transcriptional networks predicts mechanisms by which DR affects transcription via TOR.

### Tissue-specific transcriptomic effects of DR are recapitulated by TOR suppression

To assess whether the conditions of diet-induced longevity and drug-induced longevity have equivalent transcriptional effects, we first studied the equivalence of these treatments within tissues. We compared the changes in expression of all genes in the transcriptome under these two conditions, with the prediction that the induced changes would be positively correlated. As predicted, in whole flies, brains, guts, ovaries and thoraces, the diet-induced and drug-induced longevity conditions had positively correlated effects on gene expression (Figure 2). The transcriptional changes observed in these organs therefore mirror the associated changes to lifespan^10^. However, the two long-lived conditions did not have equivalent effects in the fat body, indicating that the coordination of function is largely but not obligately coupled across organs. To test explicitly the overlapping effects of DR and TOR suppression at the level of individual transcripts, differential expression analysis was performed. Surprisingly, no genes were differentially expressed in the ovary under DR, despite the robust decrease in egg laying in this condition^10^. However, in each other tissue, both lifespan-enhancing treatments had overlapping transcriptional signatures (Table 1). Overall, these analyses demonstrate that the transcriptional effects of DR largely mirror those of pharmacological TOR suppression.

**Figure 2.**
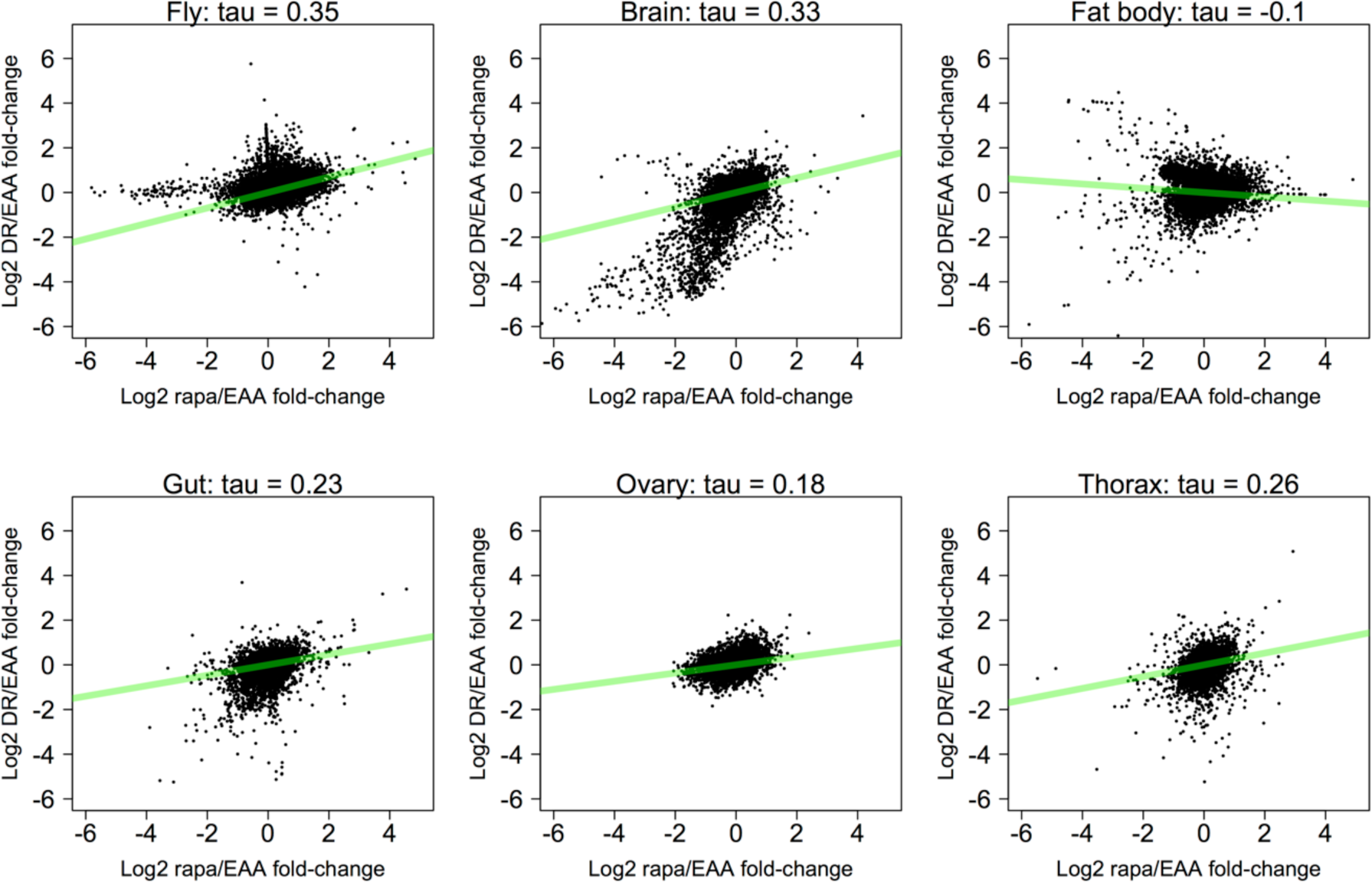
Changes in expression induced by DR correlate those induced by TOR suppression. Log2 fold-changes in expression in the diet-induced longevity and drug-induced longevity conditions, relative to the EAA-enriched control (DR/EAA fold-change and rapa/EAA fold-change, respectively), were calculated for every gene in the transcriptome. Fold-changes were calculated by DESeq2. Text above panels indicates correlations (Kendall’s Tau) between fold-changes induced by the two long-lived conditions. All p-values < 2.2e^-16^

**Table I.** Overlapping effects of the diet-induced longevity (DR) and drug-induced longevity (rapamycin) conditions on changes in gene expression within specific tissues.

| Gene set |  | Number of genes<sup>†</sup> |  |  | P-value<sup>§</sup> |
| --- | --- | --- | --- | --- | --- |
| Tissue | Expression change<sup>*</sup> | DR | Rapamycin | Overlap |  |
| Whole fly | Up | 254 | 2239 | 203 | 5.55e-121 |
|  | Down | 40 | 2159 | 17 | 3.99e-06 |
| Brain | Up | 154 | 22 | 1 | 0.02 |
|  | Down | 439 | 82 | 63 | 8.44e-82 |
| Fat body | Up | 221 | 160 | 6 | 0.02 |
|  | Down | 408 | 156 | 56 | 2.75e-46 |
| Gut | Up | 8 | 49 | 6 | 7.60e-17 |
|  | Down | 72 | 52 | 6 | 1.78e-08 |
| Ovary | Up | 0 | 30 | NA | NA |
|  | Down | 0 | 47 | NA | NA |
| Thorax | Up | 141 | 39 | 10 | 1.32e-13 |
|  | Down | 153 | 25 | 6 | 8.11e-09 |

\* Relative to EAA diet † Number of genes showing given change in expression § Hypergeometric test

To look for ubiquitous molecular signatures of lifespan extension, we examined overlaps between the transcriptional changes observed between different tissues. Surprisingly, no one transcript responded to DR/TOR in all organs, and the whole-fly samples captured only a small portion of the organ-specific changes associated with lifespan extension (Figure S1). Therefore, DR cannot be understood in terms of any one tissue. To address whether this was functionally relevant, we analysed enrichment of GO terms amongst the transcripts associated with longevity. Most GO terms (83%) were associated with specific organs, showing that there is also no ubiquitous functional signature of DR/TOR (Figure S2). Together these analyses show that the DR regulon can largely be accounted for by TOR (excepting the ovary), but that the identity of the responsive transcripts is tissue-dependent. The organ-specificity of longevity-associated changes indicates parallel responses amongst organs to EAAs and rapamycin, suggesting the coordination of organ-specific functions changes under DR and low systemic TOR.

### Diet and TOR suppression have overlapping effects on the orchestration of expression across organs

Discrete tissues compartmentalise functions, and an individual’s fitness depends on correctly orchestrating these tissue-specific functions to match the nutritional environment. It is therefore possible that the lifespan benefits of DR and TOR suppression may be mediated by certain tissues, and the biological costs by others. Therefore we investigated how DR changes the orchestration of gene expression across organs, studied as an integrated system. To identify interdependent gene expression (i.e. gene coexpression) amongst organs, we employed Weighted Gene Coexpression Network Analysis (WGCNA;^23^). This approach can identify codependent gene expression and reduce dimensionality of complex data: therefore, applying it in our study allowed us to identify codependent gene expression across organs, and changes in those codependencies in the long-lived conditions. After applying quality controls and removing genes with zero variance, 11164 of 13442 analysed genes clustered into 14 modules (Figure 3a, Figure 3b, Supplementary Materials), representing groups of genes that share expression patterns across organs. By describing modularity in our data, this network provides a complete description of the structure of the transcriptome across all organs and diets under study.

**Figure 3.**
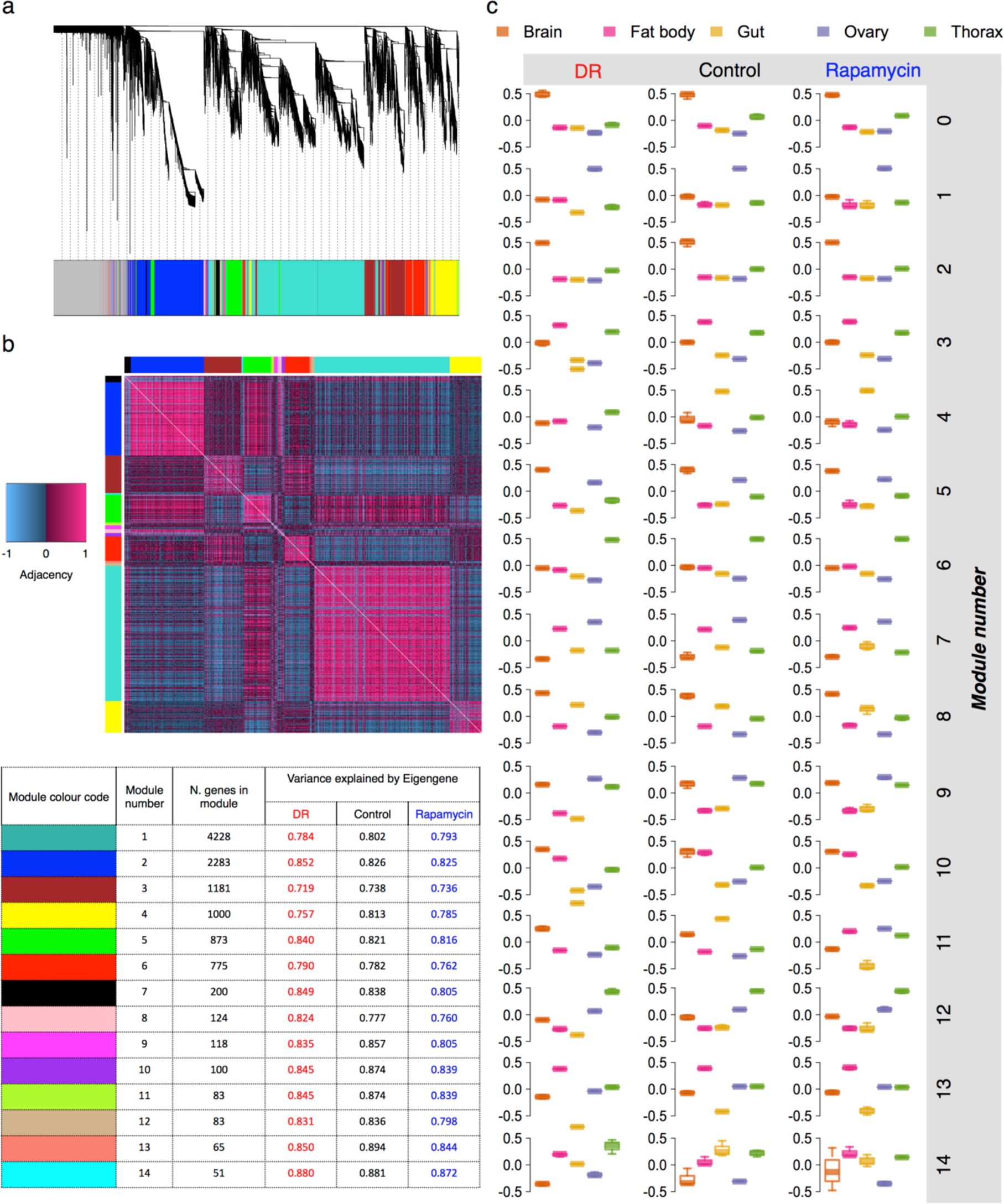
A network description of transcription across *Drosophila* organs and long-lived conditions. **(a)** Consensus transcriptional modules found across organs in all three experimental conditions. Genes were clustered according to their expression (log2 RPKM) across organs by Weighted Gene Coexpression Network Analysis (WGCNA), revealing modules of genes showing similar coexpression across organs. Leaves of the tree indicate genes. The tree was cut by hybrid tree cutting to define transcriptional modules (clusters). Tree cutting assigned genes to one of 14 consensus transcriptional modules, indicated by colour-coded vertical lines that together form the horizontal bar beneath the dendrogram (see Supplementary Spreadsheets for assignments of genes to modules). Genes not assigned to any module are colour-coded grey. **(b)** Module assignments identify clear structure in the transcriptional network, using the EAA-fed control as an example. Modules are indicated by coloured side bars. The heatmap indicates gene-gene adjacency (i.e. signed suared correlations) in expression. **(c)** Representative expression (Eigengenes) of all modules (y-axis showing Eigengene values, A.U.). These data summarise changes in transcriptional networks across organs and experimental conditions. Data are plotted by experimental condition in columns (“DR” = diet-induced longevity, “Rapamycin” = drug-induced longevity, short-lived control = “EAA”), and transcriptional module in rows. Within each plot, boxes are colour-coded by tissue (see key). Boxplots show medians (horizontal midline), 1st and 3rd quartiles (hinges), and range of data points.

We built on the transcriptional network analysis to isolate significant changes in network structure in the two long-lived conditions, by identifying pairs of modules exhibiting significantly changed coregulation. Quantifying module coregulation required reduction of the complexity of the data. Therefore, we summarised gene expression in each module with a single “Eigengene” vector per module (Figure 3c), calculated as the first principal component of expression of the genes in each module^25^. These Eigengenes accounted for between 72% and 89% of the variance in each module (weighted average = 79.86%, Figure 3b), thus simplifying the description of the organ system by two orders of magnitude, from 11.1e^3^ transcripts to 14 Eigengenes, whilst retaining ~80% of total information. We then used these Eigengenes to quantify changes in module-module coregulation. For all pairs of Eigengenes, correlations were calculated in each dietary condition then, for each long-lived condition, the correlation coefficients were subtracted from the corresponding coefficient in the EAA-fed control. This quantified changes in module-module coregulation induced in the two long-lived conditions. Repeating the same procedure on random permutations of the data generated null distributions for each pair of modules. Pairs of modules which showed significantly altered coregulation were identified by comparing the observed changes in correlation coefficients to the null distributions. This analysis thereby systematically identified pairs of modules whose coregulation was significantly altered by the conditions causing longer life (Figure 4). The diet-induced longevity condition significantly changed the coregulation of three pairs of modules (4 & 13, 4 & 3, 14 & 13). All significant changes in the drug-induced longevity condition were paired with module 11 (relative to modules 1, 4, 7, 8, 9, 12, 13). This correlation-permutation approach identified changes in coregulation of pairs of modules, but did not reveal whether those changes were driven by one or both members of each pair. Therefore, by three separate methods, we asked which individual modules’ Eigengenes were most strongly perturbed across the experimental conditions (Figure S3). These three analyses identified modules 4, 11 and 14, which were all identified by the correlation-permutation analysis. Therefore, the Eigengene analyses collectively indicated that altered regulation of modules 4, 11 and 14 changes the coordination of functions across organs in the long-lived conditions. Module 11 was associated specifically with drug-induced longevity, module 14 was associated specifically with diet-induced longevity, and module 4 was associated with both diet-induced and drug-induced longevity. These results indicate that the coexpression of module 4 with other modules may be associated generally with longevity by TOR suppression, whilst modules 11 and 14 exhibit intervention-specific changes in coexpression.

**Figure 4.**
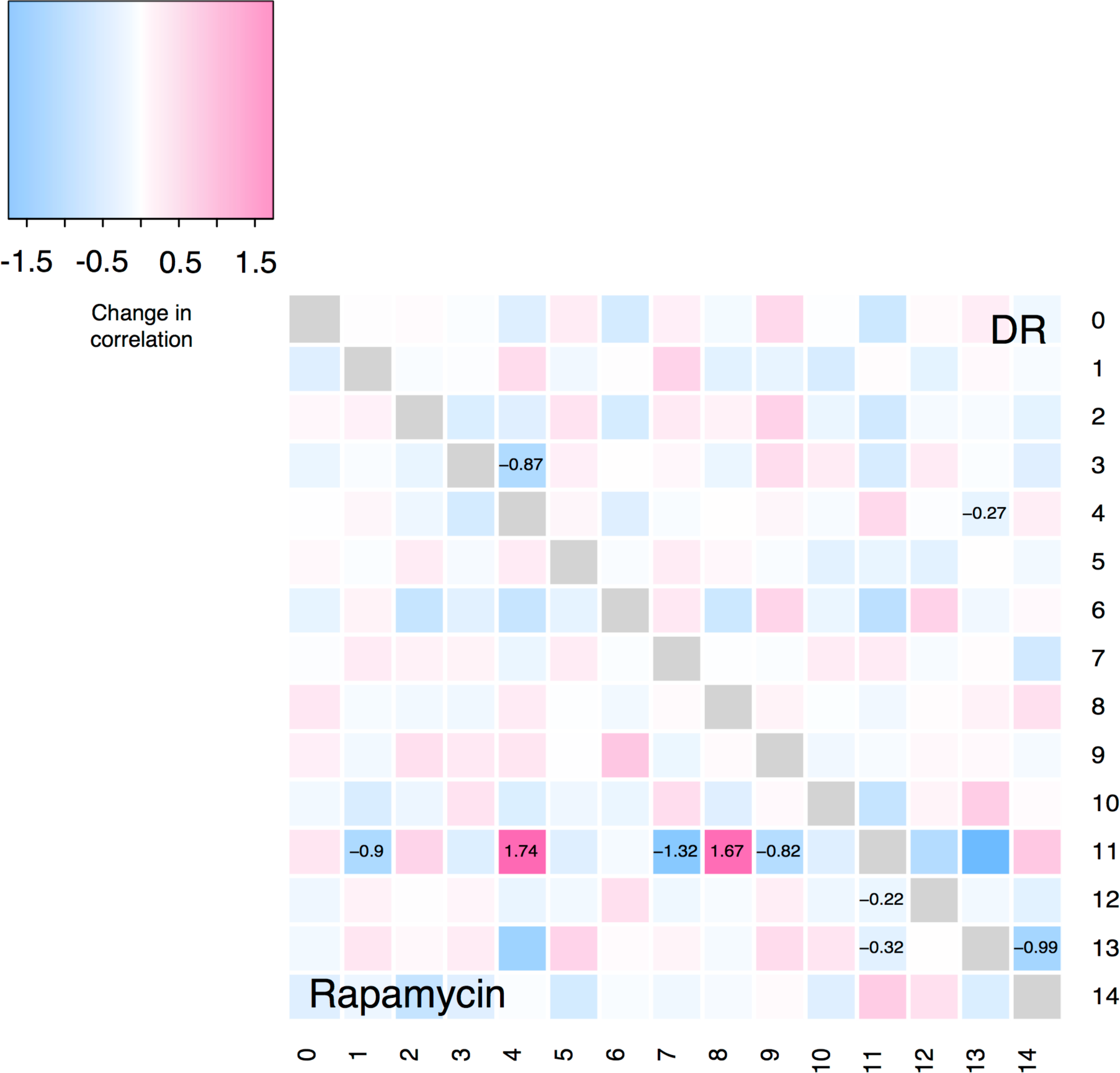
Structure of longevity-associated changes in the transcriptional network. The heatmap shows the observed change in correlation (Spearman’s rho) between pairs of modules in response to DR and in response to TOR suppression (Rapamycin), relative to the fully EAA-fed condition. Row and column labels represent transcriptional modules. Values indicate the difference in correlation between the given condition and the EAA-fed condition, and are given when the change was significant (p≤0.05) according to permutation testing.

To identify functions that are likely affected by changes to coexpression across organs, we analysed enrichment of Gene Ontology (GO) terms within each module (Supplementary Materials). Distinct transcriptional modules were associated with non-overlapping functions, indicating that these modules represent functionally distinct suites of genes, and therefore changes to their coexpression is likely physiologically relevant. Of the modules implicated in responses to DR, Module 4 was enriched in extracellular metabolic enzymes - particularly peptidases - and also lipases and carbohydrases. Module 14 was enriched in transmembrane sugar transporters and polysaccharide/carbohydrate binding. Module 11 was enriched in ATP-binding cassette (ABC) transporter activity. Taken together, the transcriptional network and GO analyses suggest that longevity is associated with altered orchestration of metabolic functions across organs, and not changes to the same metabolic program in all tissues.

### Predicting transcription factors to regulate longevity-associated transcriptional changes

We sought to identify candidate regulators of transcriptional changes associated with diet-induced longevity and drug-induced longevity. We predict that manipulating transcription factors (TFs) that regulate DR/TOR-dependent transcriptional regulation will mimic DR by correctly coordinating diverse transcriptional targets. We identified transcription factor binding site motifs (TFBSs) associated with transcriptional variation across organs in the long-lived conditions. This process associated each transcriptional module with a potentially unique set of TFs, but clustering modules by these TF sets grouped modules 4, 11 and 14 (Figure 5a), which the transcriptional network analyses had implicated in longevity. This clustering therefore suggested that these longevity-associated modules share common upstream regulators. Indeed, the sets of TFs identified in the longevity-associated modules overlapped substantially (Figure 5b), and these shared motifs were also the most significantly enriched; for each longevity-associated module, the most strongly enriched motifs bind GATA family TFs (*GATAd, GATAd, grn, pnr, srp*) (Table II). Furthermore, the most significant association observed in the entire analysis was between GATA-binding motifs and Module 4, the only module associated with both diet-induced and drug-induced longevity. Other motifs associated with all three longevity-associated modules were annotated with *Bx* - which physically binds the GATA TF *pnr*; *Ham*, which has roles in cell fate determination^26,27^; and *CG10348*, which has no known regulatory function. However, the association with these latter three transcription factors was less significant than the association with GATA factors. Looking across all modules, all but 2 and 5 were associated with some GATA motifs, however, the strength of enrichment (Escore) for the GATA factors was significantly greater for the longevity-associated modules 14 and 4 (Figure 5c). The final evidence of association between GATA factors and longevity-associated modules was that Module 4 was not only highly enriched in GATA binding sites, but also contained GATAe (Supplementary Files). Experimental studies have shown correlated expression of a TF and its putative targets is a strong predictor of causal relationships^21,22^. Altogether, these findings strongly implicated GATA factors in the tissue-specific transcriptional changes observed under low TOR.

**Figure 5.**
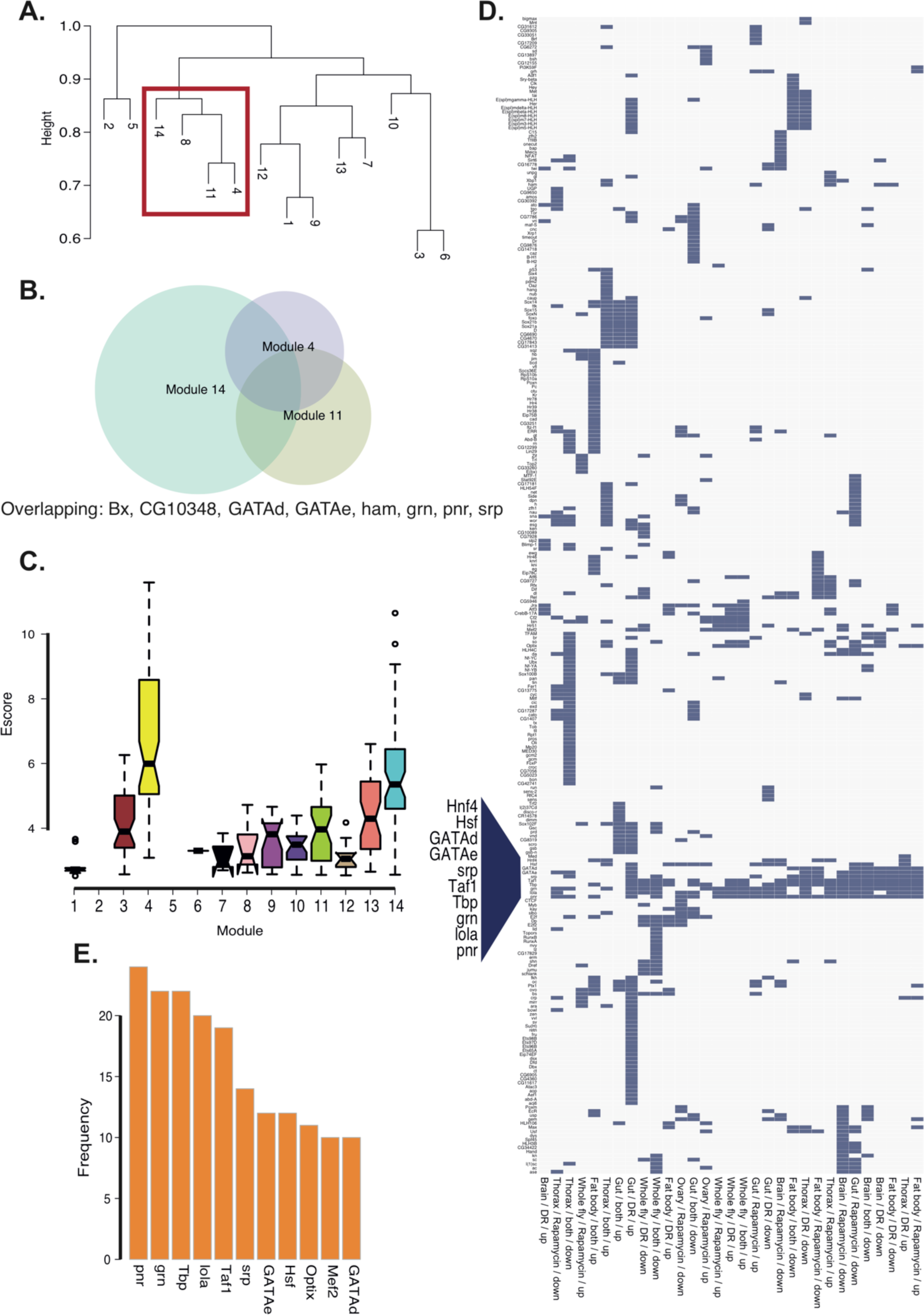
A signature of GATA transcription factors as regulators of the transcriptional response to DR and/or TOR. **(a)** Lifespan-associated transcriptional modules are grouped by sharing of transcription factor binding site motifs. Each module was tested for enrichment of *cis*-regulatory elements, and clustered by the presence/absence of associated transcription factor binding site motifs. Dendrogram labels represent module numbers. This analysis clusters together transcriptional modules associated with longevity (4, 11, 14, also clustered with module 8). **(b)** Longevity-associated modules are associated with overlapping sets of transcription factor binding sites. Text below the Venn diagram names the transcription factors associated with binding sites enriched in all three longevity-associated modules. **(c)** Distribution per module of enrichment (Escore) of motifs annotated as binding GATA transcription factors. The box plots show medians (horizontal midline), 1st and 3rd quartiles (hinges), and range of data points. The notches equate to ~95% confidence intervals for the medians [(1.58 * interquartile range) / (square root *n*)]. The plot shows that enrichment of GATA binding sites is particularly strong for modules 4 and 14, both of which are associated with lifespan extension by DR. **(d)** GATA factor binding sites are enriched across multiple gene sets regulated by DR/TOR. Columns represent sets of genes according to tissue and the sign of expression change per treatment. Rows represent TFs associated with the sets of differentially expressed genes by enrichment of binding site motifs in the gene set. The axes are ordered by clustering the presence (blue) and absence (grey) of TF binding sites, using binary distance. Names of transcription factors occurring in a cluster across multiple gene sets have been magnified. **(e)** The bar plot shows the frequency of associations with longevity-associated transcriptional changes within organs, for the most frequently associated TFs. TFs associated with <10 gene sets are excluded.

**Table II.**
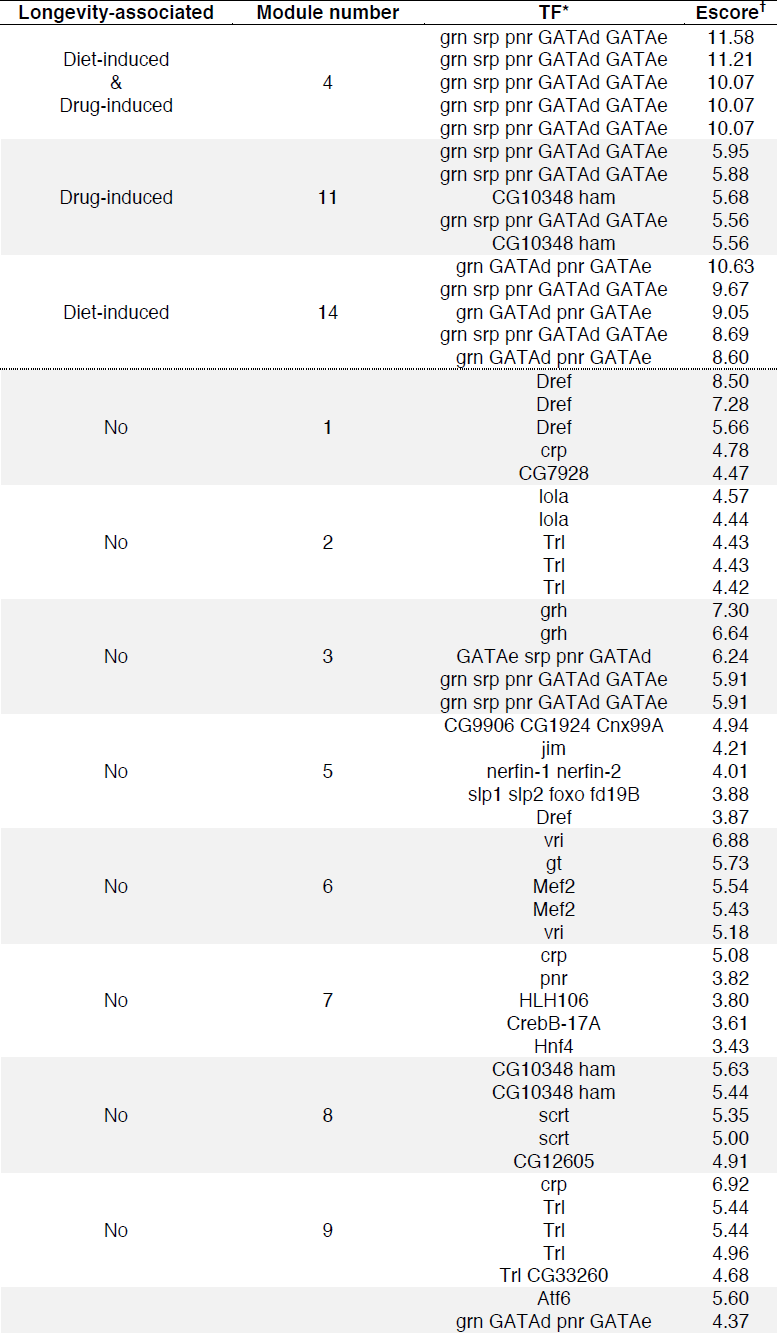
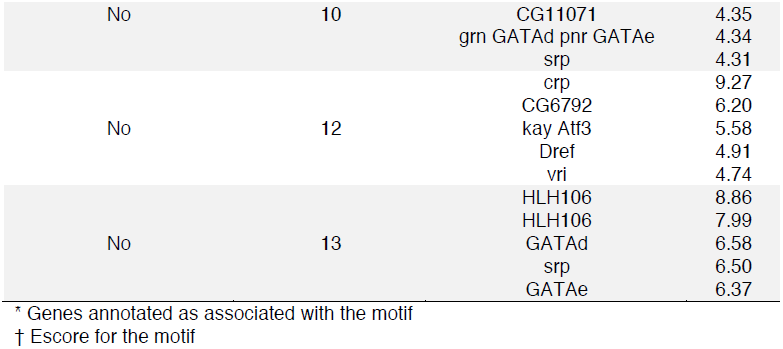
Annotations for top five-ranked motifs enriched in association with transcriptional modules. Unannotated motifs are excluded.

To complement the analysis of transcriptional network regulators, we repeated the same approach on sets of genes which showed changed regulation (differential expression) within organs in one or both of the long-lived conditions (Table III). Consistent with the absence of an organ-nonspecific transcriptional signature of DR/TOR, most TFBSs identified by this analysis were associated with specific tissues. However, one cluster of TFs was associated with multiple gene sets across organs, and that cluster contained the GATA factors (Figure 5d). Across all gene sets, each GATA factor was associated with at least 32% of diet/tissue-specific gene sets (Figure 5e). GATA factors were therefore associated with DR/TOR-dependent transcriptional changess within organs, recapitulating the parallel observations across organs.

**Table III.**
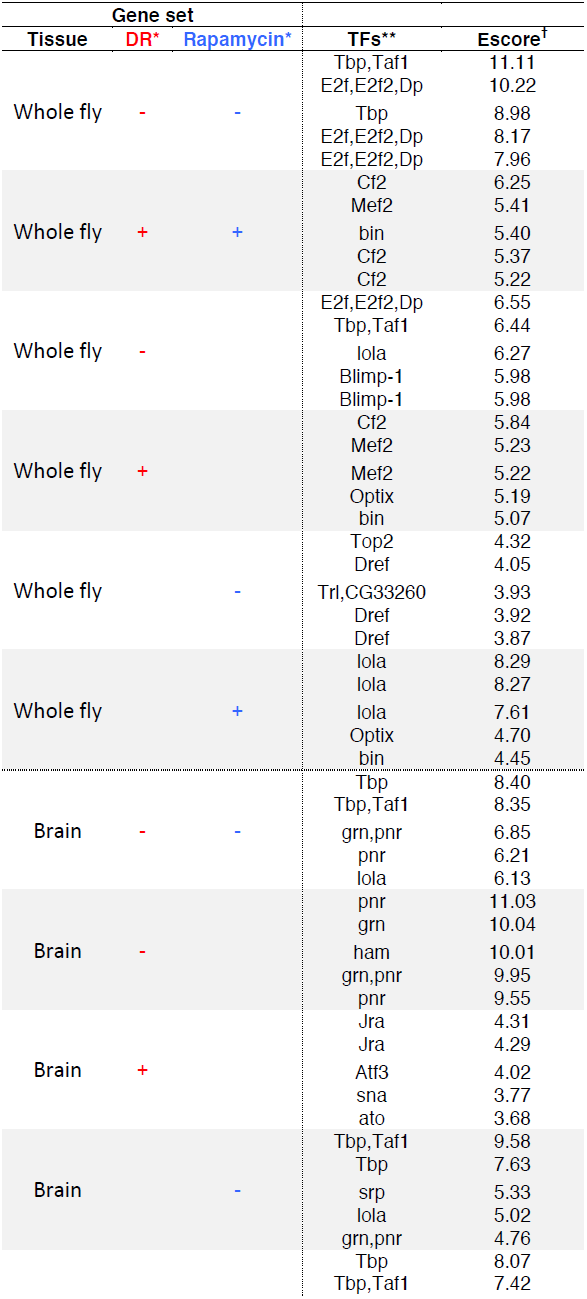
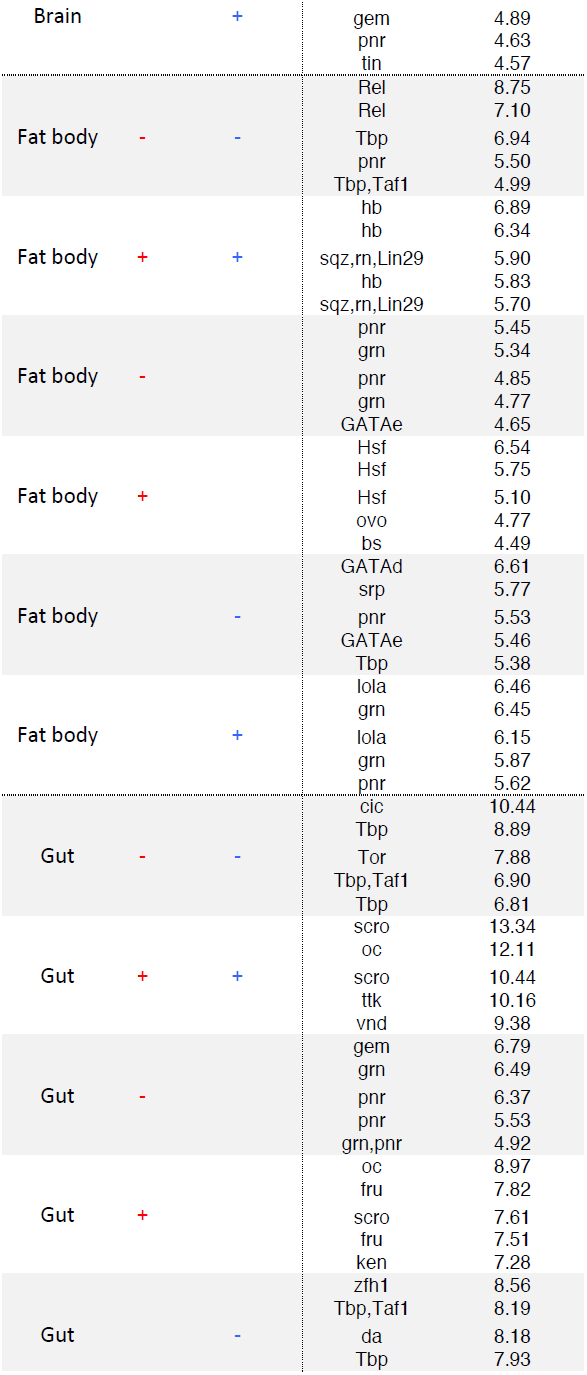
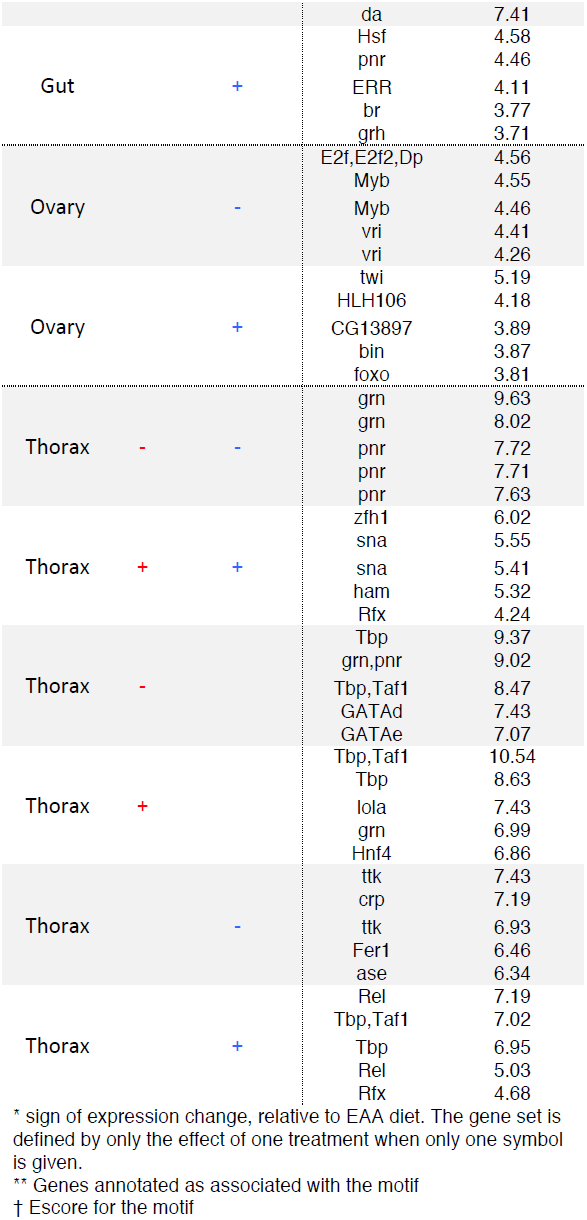
Annotations for top five ranked motifs enriched in association with differentially expressed gene sets. Unannotated motifs are excluded.

The TFBS enrichment analysis associated GATA factors both with transcriptional changes within organs and altered functions of the organs as a collective whole. This association was consistent with prior knowledge of the evolutionarily conserved biology of these TFs. GATA factors play known roles in signalling amino acid availability via TOR in evolutionarily diverse eukaryotes (e.g. yeast:^28^; and mosquitos:^29^), as well as being required for lifespan extension by some tissue-specific genetic lesions in worms^20^. To test for evolutionarily conserved connections between GATA factors and DR-responsive transcripts, we established the association between TFBSs and orthologs of DR-responsive genes across 12 *Drosophila* species. These 12 *Drosophila* species span ~40 of million years of evolution, which due to their rapid development time, equates to a greater evolutionary time than that for the mammalian radiation. The method used for this analysis was independent of that used for *D. melanogaster*, but also showed that the association between GATA motifs and orthologs of DR-responsive genes from *D. melanogaster* was robust across *Drosophila* species (Supplementary Spreadsheets). Together, these data and previous findings strongly implicate GATA factors as evolutionarily-conserved regulators of tissue-specific transcriptional responses to EAAs.

### Functional roles for GATA factors in mediating dietary effects on lifespan

Of the five *D. melanogaster* GATA TFs, *GATAe* and *srp* were of particular interest for a role in longevity via DR/TOR. *GATAe* was associated more strongly than any other TF with longevity-associated transcriptional changes (see above), and *srp* has known roles in the fat body in regulating oogenesis via yolk proteins^30^. Consequently we focussed on testing the roles of *srp* and *GATAe* in dietary regulation of longevity. We knocked down *srp* and *GATAe* with tissue-specific RNAi, and evaluated whether these knockdowns altered the effects of dietary EAAs on egg laying and lifespan. *GATAe* and *srp* were most highly expressed in the gut and fat body, respectively (Figure S4), so we targeted *GATAe* in the midgut epithelium (with the TiGS driver:^31^) and *srp* in the fat body (with the S1106 driver, which drives in both fat body and gut^32^, however fat body-specific expression of *srp* (Supplementary Materials) suggests that RNAi phenotypes can be attributed solely to the fat body). These GeneSwitch drivers^33^ are activated by feeding flies the RU_486_ inducer (RU).

In *GATAe^RNAi^* flies there was a significant interaction between the effect of feeding the transgene-inducing drug and EAA on the lifespan of GATAe knockdown flies (Cox Proportional Hazards Regression, P < 0.005). Specifically, knocking down *GATAe* in the gut accelerated the onset of mortality in early life but, surprisingly, this lifespan shortening was partially rescued by feeding dietary EAAs (Figure 6a). However GATAe^RNAi^ affected neither egg laying nor altered the effect of EAAs on egg laying (Figure 6b), indicating that the regulation of egg laying is independent of the lifespan-limiting process induced by knocking down GATAe in the gut.

**Figure 6.**
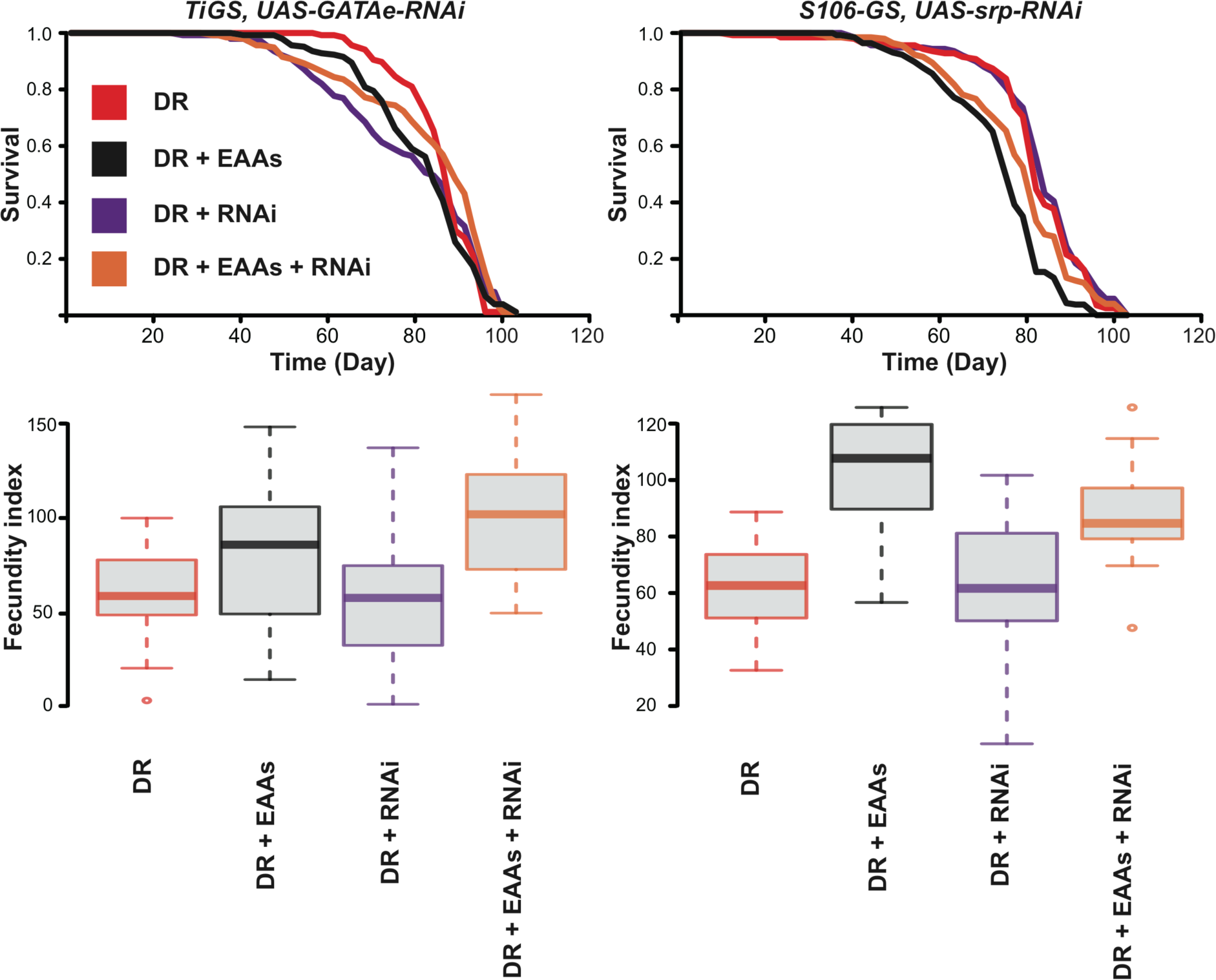
Tissue-restricted knockdown of *GATAe* and *srp* interact with dietary EAAs to determine egg laying and lifespan. Survival curves **(A & C)** and egg laying indices (overnight egg laying by vials of 10 flies; **B & D**) of flies expressing *GATAe^RNAi^* in the gut (A, B), or *srp^RNAi^* in the gut and fat body.

In contrast to *GATAe* knockdown, *srp* knockdown had positive effects on longevity (Figure 6c). Fat body-specific expression of *srp^RNAi^* protected flies from the lifespan-shortening effects of EAAs (log-rank test, p<0.001), but there was no effect on lifespan in the absence of EAAs (p=0.57). Egg laying was not affected by srp knockdown (p=0.08). Therefore, knocking down *srp* in the fat body insulates lifespan against the pernicious effects of EAAs, but does not preclude flies from the early-life benefits of EAA feeding for egg laying. Together, the GATAe^RNAi^ and srp^RNAi^ experiments validate our bioinformatic analysis, by linking age-dependent physiology with interactions between dietary EAAs and the tissue-specific regulation of GATA factors.

Our expression data showed that *srp* is expressed most strongly in the fat body, but is also expressed in other tissues e.g. the ovary. Therefore, we asked whether systemic *srp* knockdown using a *Daughterless* GeneSwitch driver further altered fly phenotype. Ubiquitous knockdown of *srp* was sufficient to neutralise the lifespan-shortening effects of EAAs, restoring lifespan to that of flies fed DR food (Figure 7a). Furthermore, ubiquitous *srp* RNAi extended lifespan of flies on DR food, meaning that transgene expression resulted in a median 7 day lifespan extension whether or not the flies were fed high levels of dietary EAAs. Whilst systemic *srp* knockdown extended lifespan more strongly than the tissue-specific knockdown, the tradeoff with egg laying was much greater (Figure 7b). Without induction of *srp^RNAi^*, the flies laid ~25% more eggs when EAAs were supplemented to the food, but *srp* knockdown arrested egg laying (ANOVA, p < 2.2^-16^) and rendered it insensitive to EAA feeding (EAA * RU interaction, p<0.005). The egg laying phenotype thus contrasted the lifespan phenotype, since egg laying was entirely desensitised to diet, whereas median lifespan remained sensitive. Together, these data indicate that the relationship between lifespan and egg laying is set by an interaction of dietary EAAs and *srp*, and the integration of egg laying and lifespan depends on the tissues in which this interaction occurs.

**Figure 7.**
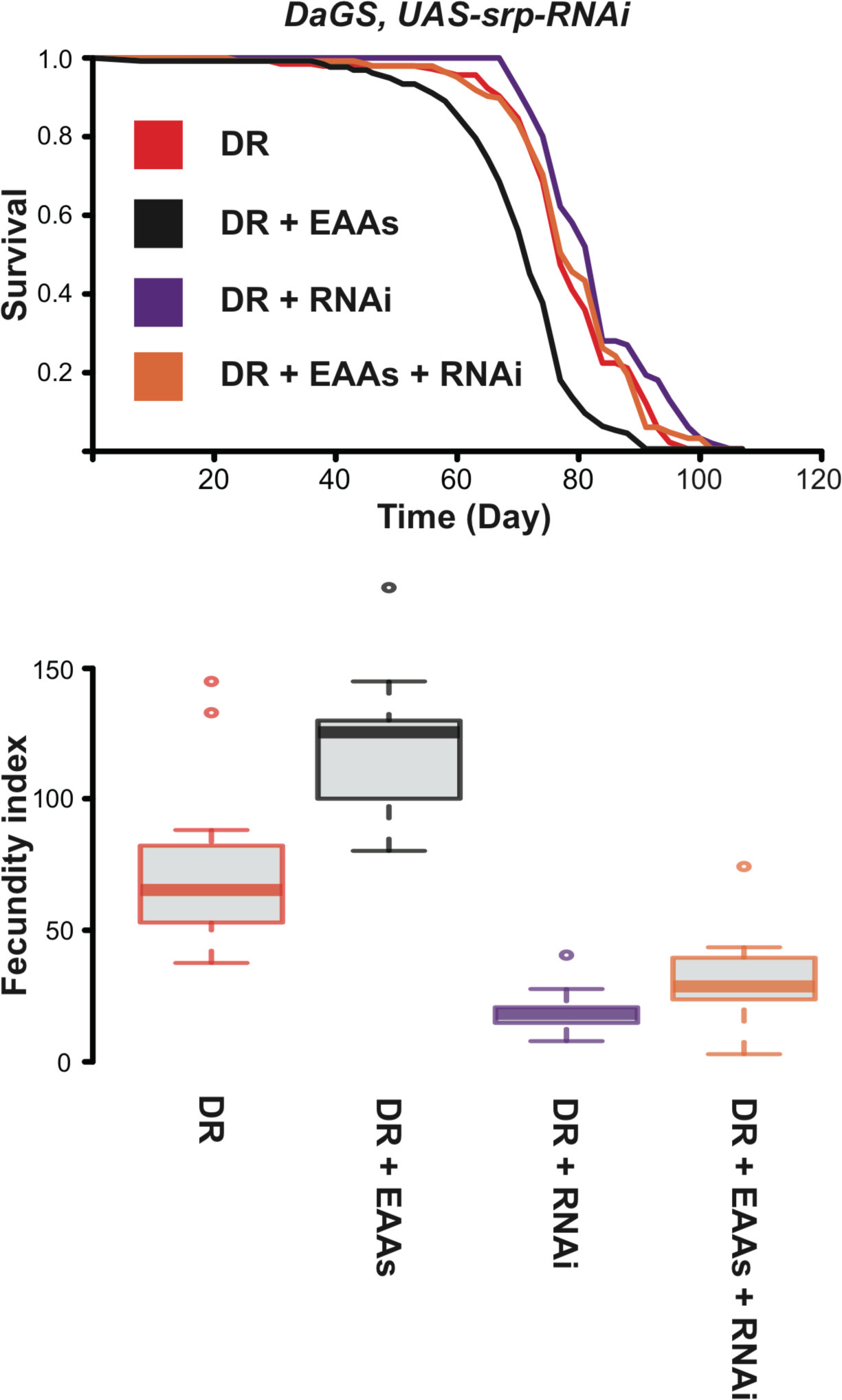
Ubiquitous knockdown of *srp* reverses lifespan-shortening effects of EAAs and desensitises egg laying to EAAs. **(A)** Survival curves and **(B)** egg laying indices of flies ubiquitously expressing RNAi against *srp*.

## Discussion

Dietary restriction improves lifelong health and extends lifespan in a range of organisms, from yeast to mammals. A growing body of evidence shows the particular importance of dietary nutrient balance for ageing, particularly lowered protein:carbohydrate ratio^6,34^, which implicates amino-acid sensitive TOR signalling^11^. This implicit role of TOR was recently validated in *Drosophila*, by the demonstration that active TOR is required for EAA enrichment to shorten lifespan^10^. These results have the important implication that the lifespan effects of EAA restriction are not accounted for solely by metabolic effects, but also by cellular signalling. Using the same experimental conditions, our present study predicts lifespan-relevant and EAA-dependent signalling mechanisms, by showing that EAA restriction has transcriptional effects which are congruent - both within and between organs - with those of pharmacological TOR suppression, indicating that the transcriptional effects of DR are indeed mediated in large part by TOR. Furthermore, these overlapping transcriptional changes are also associated with overlapping sets of *cis* regulatory elements. Thus, DR and rapamycin have overlapping effects on lifespan, on transcription within organs, on the coordination of functions across organs, and their transcriptional targets share predicted regulators.

Our results connect lifespan regulation by DR and low TOR with GATA transcription factors. The GATA factor knockdown experiments validate the bioinformatic connections between dietary EAAs, transcription factors and lifelong physiology. Specifically, these experiments show that genetic knockdown of either *GATAe* or *srp* substantially alters the effect of diet on fly survival and, in the case of srp, knockdown both extended lifespan and dictated the influence of diet on egg laying, thus phenocopying flies treated with rapamycin. Curiously, *GATAe* knockdown in the gut changed the sign of the effect of EAAs on lifespan, suggesting either a different optimal dietary balance in these flies, or that elevated EAAs rescued the pathological effects of GATA knockdown. By contrast, *srp* knockdown with two different drivers cancelled the lifespan-shortening effects of EAA enrichment. Furthermore the GATA knockdown experiments show that, whilst lifespan and egg laying normally show coupled responses to diet, they can be decoupled by genetic manipulation of GATA factors, depending on which tissues are targeted. The phenotypic differences between flies in which srp is locally or systemically knocked down provide key evidence that tissue-targeted manipulation of TFs associated with DR can recapitulate the lifespan benefits of DR, and that the associated costs depend on the tissue-specificity of the intervention. This supports the notion that ageing can be ameliorated by specifically targeting pathologies in lifespan-limiting tissues. Whilst there is some existing evidence of a role for GATA factors in lifespan regulation^20^, to our knowledge they have not yet been associated explicitly with DR, nor has there been a prior demonstration of any transcriptional intervention blocking the phenotypic effects of EAAs. The indication that costs and benefits of dietary alteration are mediated by distinct tissues has the important biomedical implication that lifelong health may be improved by altering the coordination of function across multiple tissues, and the evolutionary implication that phenotypic tradeoffs evolve due to coupled functions of tissues.

The GATA factors are an ancient family of transcription factors, which have well-characterised and essential roles in coordinating development and growth in organisms from yeast to mice. In multicellular differentiated organisms, GATA factors are required in the development of multiple tissue types, which in *Drosophila* includes the heart^35^, fat body^36^ and gut^37^. GATA factors are also expressed tissue-specifically in adulthood (Figure S4), but their roles in coordinating adult functions are not well characterised. In *Drosophila*, recent studies have shown a role of *GATAe* in maintenance of cell identity and tissue function in the adult gut, but it is not yet clear from which molecular pathways *GATAe* integrates information^38–40^. One of the better-described roles for GATA factors in adult animals is in nutrient regulation of oogenesis in mosquitos^29,41–43^. In *Aedes aegypti*, oogenesis requires a blood meal, which contains the mosquito’s only source of protein. Egg production is suppressed before feeding, due in part to GATA-mediated repression in the fat body of the major yolk precursor protein gene *Vg*^44^. After the blood meal, *Vg* expression is activated by TOR enhancing expression of the transcriptional activator *AaGATAa*^43^. In *Drosophila*, there is evidence that this regulatory circuit may be conserved, since *srp* regulates expression of yolk proteins^30^. Our new and recent data show that rapamycin abrogates higher egg laying caused by EAA enrichment^10^, corresponding to changes in the fat body in expression of yolk proteins (Supplementary Materials). The connection in *Drosophila* between dietary nitrogen and GATA transcriptional control is consistent with mechanisms in the yeast *Saccharomyces cerevisiae*, in which selective amino acid catabolism is controlled by a circuit known as Nitrogen Catabolite Repression. In this system, when the available nitrogen sources only support poor growth, TOR-dependent nuclear localisation of a GATA transcription factor triggers the expression of genes involved in the transport and metabolism of less-preferred nitrogen sources^28^. Together, these data point to highly evolutionarily conserved connections between protein uptake, growth and reproduction, TOR signaling and transcriptional control by GATA factors. Theory suggests that ageing results from antagonistic pleiotropic effects of mechanisms that promote growth and reproduction in the young^45^. GATA factors fit these criteria, as nutrient-responsive regulators of growth that we associate with molecular responses to lifespan-extending regimes. Due to the evolutionarily conserved connections between dietary nitrogen and lifespan on one hand, and dietary nitrogen and GATA factor-regulated transcription on the other, we anticipate new interest in the role that GATA factors play in lifespan regulation.

Our experiments demonstrate that the tissue-restricted knockdown of GATA factors is sufficient to modify the lifespan-shortening effect of EAAs. Previous studies have shown that tissue-targeted interventions to lower insulin signalling are sufficient to extend lifespan in worms and flies^46^, and GATA factors are required for some such effects. It is well-established that the balance of dietary nutrients has the evolutionarily conserved capacity to determine lifelong health^11^, and to recognise this balanced supply, insulin and TOR signalling must coordinate. Reducing either insulin or TOR signalling is sufficient to extend lifespan, suggesting that the lifespan-extending effects of these two interventions may be mediated by the nexus of their signalling effects. In *C. elegans*, the GATA factor *ELT-2* is required for longevity following dietary restriction or mutation of the insulin receptor^20^, and GATA factor overexpression extends lifespan^47^. Additionally, previous transcriptional studies in both flies and worms have uncovered enrichment of GATA motifs in the insulin regulon^48,49^. Together with our data, these results suggest that signals from TOR and other nutrient sensing pathways are mediated at least in part by GATA factors. In light of these data, our demonstration that transcriptional effects of diet are largely tissue-specific suggests that TOR and the GATA factors mediate cell-autonomous interpretations of global signals (e.g. insulins or bioamines), into a local language that dictates physiological change appropriate to the tissue in question.

Transcription factors known to be downstream of TOR signalling and insulin signalling are candidates for interpretation by GATA factors in *Drosophila. REPTOR* and *REPTOR-BP* have recently been discovered by Tiebe and colleagues as a novel TOR-dependent transcription factor complex^18^. Given the complementary nature of Tiebe *et al*’s cell culture experiments and our study of dissected tissues, an emergent hypothesis is that GATA factors determine organ-specific responses to TOR by modifying effects of systemic *REPTOR* signalling. Downstream of insulin signalling, molecular interactions between *FoxO* and TOR signalling are already established: TOR lies in a network of interacting signaling pathways including insulin^50^, and *FoxO* is required for reduced insulin signalling to extend lifespan^51^. However, FoxO binds *TOR*’s promoter and is required for normal *TOR* expression, and GATA motifs are associated with genes that are not bound by *FoxO* but show *FoxO*-dependent regulation by IIS^49^, suggesting a regulatory circuit from IIS to GATA factors via *FoxO* and TOR. Given that *FoxO* is not required for the lifespan-extending effects of DR^52^ and that different insulin-like peptides are produced in response to different nutritional stimuli^53^, we suggest that signals from IIS via FoxO may be modified in peripheral tissues by TOR-modulated GATA activity, which shapes tissue-specific responses to diet that ultimately modify physiology and longevity (Figure 8).

**Figure 8.**
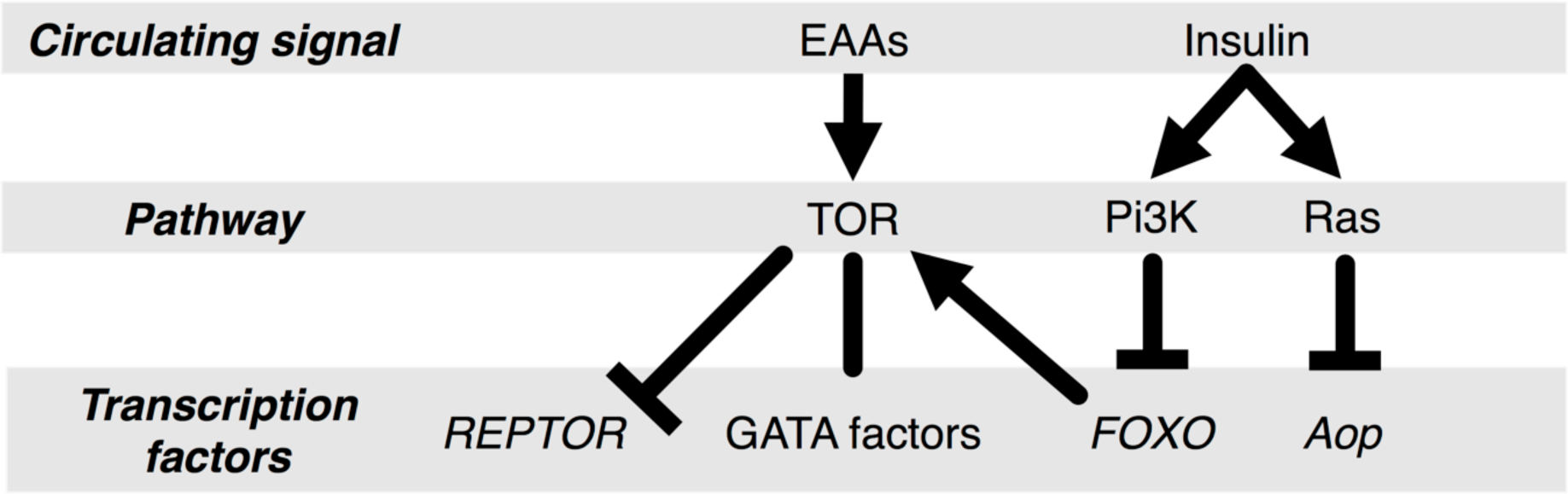
Predicted molecular integration of GATA factors in nutrient-sensing and lifespan-relevant transcriptional networks in *Drosophila*. Our transcriptomic analysis associates transcriptional effects of EAA restriction with TOR, and the TOR-dependent EAA regulon with GATA factors. Previous studies have shown that (1) TOR directly represses *REPTOR*, (2) The insulin pathway bifurcates via Pi3K and Ras to regulate lifespan via *FOXO* and *Aop*, and (3) GATA binding sites are associated with genes that are differentially expressed in *FOXO* mutants but not bound by *FOXO*, that *FOXO* both binds *TOR*’s promoter and *TOR* transcription is reduced in *FOXO* mutants. Together, these results suggest that *TOR* integrates signals from IIS/Pi3K signalling via *FOXO*, that regulons of DR, TOR and *FOXO* are all enriched in GATA factor binding sites, and therefore that GATA factors mediate transcriptional effects of both EAAs and *FOXO* via TOR.

The gut appears to be a particularly effective target for interventions to extend lifespan^46^. The adult gut has a critical role in the ongoing health of organisms, balancing the passage of nutrients whilst resisting environmental stresses^54^. In flies, there are complex relationships between age, gut maintenance, diet, metabolism, resident microbiota and expression of antimicrobial genes^55–62^. Recent work suggests a central role for the gut in mediating lifespan extension by DR in female flies, revealing sexually dimorphic pathologies in aged female guts^63^, dependent on cell-autonomous expression of sex determination pathways^64^. Failure of gut integrity is associated with changes in microbiota and greater antimicrobial peptide expression, and appears to be a marker of imminent death^58,65^. In worms, the GATA factor ELT-2 interacts with p38 transcriptional regulators to modify adult gut immunity^66^, whilst GATA factors are required for normal gut development and maintenance in *Drosophila* and mice^38,67^. It is thus tempting to speculate that increased risk of death in late life may be preciptated by a loss of gut integrity enhancing exposure to environmental microbes and toxins. Our results are directly relevant to these issues, because coordination across organs of a transcriptional module (module 4) that was perturbed under long-lived conditions strongly corresponded to *GATAe* expression, and *GATAe* and paracrine insulin signalling have each been shown to be required for intestinal stem cell proliferation^38,68^. Our lifespan data are consistent with *GATAe* knockdown in the gut increasing mortality, but since this effect appears to be stochastic and not necessarily connected to age, the role of *GATAe* regulation in late-life mortality remains to be established. However, fly lifespan is not necessarily extended by enhancing expression of components of the immune system^69^, antibiotic treatment^55^ or genetically reducing gut dysplasia^59^. Thus, although compromised gut integrity is clearly a marker of frailty in late life, it may not be obligately linked to death.

This study reveals how the fly’s tissues interact as a system, and how that system responds to EAA dilution and low TOR. The results showed that the effects of EAA restriction on transcription are likely mediated by TOR via GATA transcription factors. Entirely consistent with this prediction, genetic analysis confirmed that the tissue-specific activity of GATA factors dictate the effect of dietary nutrient balance on phenotype. Importantly, these experiments also suggest that the costs and benefits of dietary variation may be mediated by different tissues, and therefore that benefits may be reaped without fitness tradeoffs by targeting specific tissues. The evolutionary conservation of GATA factors, of their connection to regulating amino acid metabolism, and of the capacity of TOR to mediate lifespan extension, suggests that GATA factors may be relevant to ameliorating ageing by DR in a broad range of organisms, including humans.

## Materials & methods

*Drosophila melanogaster* and diets were prepared according to (10). 1SY medium is a dietary restriction (DR: diet-induced longevity) regime that extends lifespan in wild-type and laboratory-maintained flies^71,72^, containing 100 g/l autolysed yeast (MP Biomedicals, OH, USA), 50 g/l sucrose (Tate & Lyle, London, UK), 15 g/l agar (Sigma-Aldrich, Dorset, UK), 30 ml/l nipagin (Chemlink Specialities, Dorset, UK), and 3 ml/l propionic acid (Sigma-Aldrich, Dorset, UK). EAA food comprised 1SY with the addition of cocktail of EAAs dissolved in pH 4.5 water (final concentrations in fly media: L-arginine 0.43 g/l, L-histidine 0.21 g/l, L-isoleucine 0.34 g/l, L-leucine 0.48 g/l, L-lysine 0.52 g/l, L-methionine 0.1 g/l, L-phenylalanine 0.26 g/l, L-threonine 0.37 g/l, L-tryptophan 0.09 g/l, L-valine 0.4 g/l: all suppled by Sigma). EAA+rapamycin (drug-induced longevity) food consisted of EAA food with the addition of rapamycin (LC laboratories, MA, USA) dissolved in ethanol, to a final concentration of 200 μM in the diet. For RNAi experiments, RU_486_ (Sigma M8046) dissolved in ethanol was added to 1SY or EAA food to a final concentration of 200 μm. Vehicle controls were added to media as appropriate.

Outbred wild-type Dahomey flies bearing the endosymbiont *Wolbachia* were cultured on a 12:12 light cycle at 25oC and 60% humidity. For RNAi experiments, the TiGS and S1106 drivers and RNAi constructs (UAS-srp^RNAi^: Vienna Stock Center #33748; UAS-GATAeTRiP: Bloomington Stock Center #33748) were backcrossed into flies bearing the *w* mutation for at least 6 generations (UAS-GATAeTRiP was back-crossed by PCR), and maintained at large population sizes to maintain outbred genetic diversity. For all experiments, parents of experimental flies oviposited onto grape juice agar for 18h. Eggs were washed from this agar, added to 1SY and cultured to adulthood at standardised density. Newly emerged flies were allowed to mate *ad libitum* for 48h before being lightly anaesthetised with CO2. Males were removed, and female flies were allocated to experimental diets. For RNA sequencing, females were maintained on 1SY, EAA or EAA+rapamycin for six days before dissection. The RNA sequencing experiment was independently replicated three times, generating three samples per organ or per whole fly, per diet (with the exception of the gut on 1SY: see Supplementary Text). For egg laying and lifespan experiments, survival was scored three times per week. Egg laying was scored on day 10 after eclosion, after 18h egg laying. Fecundity indices were calculated as number of eggs per female (n=10 females per vial).

RNA was collected 6-10h into the flies’ light cycle. To prepare RNA for sequencing, whole flies were flash-frozen. Brains, abdominal fat bodies, ovaries, guts and thoraces were micro-dissected in ice-cold RNAlater solution and frozen at −80°C. RNA was extracted using the QIAGEN total RNA isolation kit and quantified on an Agilent 2100 bioanalyser. Sequencing was performed by the high throughput genomics services center at the Huntsman Cancer Institute (University of Utah). Sample concentration and purity of RNA was measured on a NanoDrop spectrophotometer, and RNA integrity was assessed on an Agilent 2200 TapeStation. Illumina TruSeq libraries were prepared from this RNA with the Illumina TruSeq Stranded mRNA Sample Prep kit and sequenced on an Illumina HiSeq2000 101 v3 platform using paired-end sequencing.

Reads were aligned to the *D. melanogaster* genome annotation 5.57 using TopHat2 2.0.14 and counted using HTSeq 0.5.4p3^73,74^. Non-protein coding genes were retained. Unmapped reads were discarded. Enumerated reads were then analysed in R (3.0 & 3.1) using BioConductor. RPKM was calculated from read counts generated by HTSeq, using the EdgeR library.

Differential expression across the three experimental conditions was determined with a negative binomial GLM fitted by DESeq2 (1.8.1,^75^), without rejection based on Cook’s distance, calculating P-values with a two-sided Wald test, and calculating false discovery rate by Storey’s method. Intersections between gene sets and enriched GO terms were visualised with the upset package in R^76^. GO term enrichent was analysed using GOrilla (http://cbl-gorilla.cs.technion.ac.il/).

Unsigned gene coexpression networks were determined using data from all organs, excluding the whole-fly samples, using the WGCNA package in R^23^. Consensus modules were determined automatically using the *blockwiseConsensusModules* function with default settings and a power of 26, stipulating a minimum module size of 50 genes. Eigengene determination, variance explained by Eigengenes and clustering were performed using internal WGCNA functions, as per the package tutorial (http://labs.genetics.ucla.edu/horvath/CoexpressionNetwork/Rpackages/WGCNA/Tutorials/index.html). Changes in between-module correlations by experimental condition were calculated by custom R functions: correlation matrices (Spearman’s rho) for module Eigengenes were calculated in each experimental condition, and changes were calculated by subtracting the DR matrix and the EAA+rapamycin matrix from the EAA matrix, to calculate observed changes in correlations between pairs of Eigengenes under DR and under rapamycin administration, respectively. To generate null distributions for changes in correlations for each pair of modules in each of the two long-lived conditions, the same procedure was repeated 10,000 times, permuting each Eigengene and calculating changes in correlation for each pair. Observed changes were considered significant when they did not fall between the 2.5th and 97.5th percentiles of their respective null distribution.

The most likely acyclic network between modules was determined by Additive Bayesian Network analysis of module Eigengenes using the R package ABN with 10000 iterations. This approach determines the likelihood of inter-dependencies between variables by randomly simulating data and comparing to the observed interdependencies. ABN found a consensus structure to the data after only ~1000 iterations, indicating that the structure is robust to further simulation. The ABN was plotted in Cytoscape, using hierarchical network ordering. Meta-module analysis and Eigengene perturbation analysis were performed according to^25^.

Enrichment of cis-regulatory motifs was analysed using i-Cis target^77^, excluding genes for which DEseq2 models did not converge. Unannotated CRMs were excluded from further analysis. Samples were hierarchically clustered according to the presence/absence of transcription factor binding sites with the R hclust function, using a binary distance metric. Notches on boxplots, approximating 95% confidence intervals of medians were produced using the “notch” argument to the R boxplot function. Heatmaps of TF:gene-set associations were plotted using the heatmap.2 function from the R gplots package, and ordered by heirarchical clustering using binary distance.

Analysis of evolutionary conservation of the association between TF binding motifs and differentially expressed genes was conducted per tissue. Groups of differentially expressed genes (according to DEseq2 analysis) showing the same signs of fold-change in response to EAA restriction were scanned for significantly overrepresented motifs in their promoter regions. 706 motifs were included in the search, from OnTheFly (2014 release), flyreg v2, dmmpmm2009 and idmmpmm2009 databases, all provided by MEME 4.10 (http://meme-suite.org). Pscan version 1.2.2 was used to score the motifs for the gene list, using the sets of genes that either didn’t show any diet-induced differential expression in any tissue, or were not expressed, as a ‘null hypothesis’ background. For each gene list, the 706 scores (one per motif) produced by pscan were normalized to *z* scores, and *P* values were computed, corrected for multiple testing (Benjamini-Hochberg).

The same procedure was repeated for the evolutionarily conserved regulation analysis, except that the genes considered were obtained from the 12 *Drosophila* species (*D.melanogaster* and *D.ananassae*, *D.erecta*, *D.grimshawi*, *D.mojavensis*, *D.persimilis*, *D.pseudoobscura*, *D.sechellia*, *D.simulans*, *D.virilis*, *D.willistoni* & *D.yakuba*) whose genomic sequences are part of FlyBase, taking only 1:1 orthologs of genes which DESeq2 analysis had shown to be differentially expressed in *D. melanogaster*. Gene selection was done in two steps: for all gene groupings (differential expression and background), genes for which OrthoDB version 7 didn’t predict a one-to-one ortholog in all other 11 species were removed. Then, each grouping was expanded with all the relevant orthologs predicted by OrthoDB, such that each gene group contained 12 times as many genes at after the first step.

Survival data were analysed in Microsoft Excel for log-rank statistics, or in R for Cox Proportional Hazards analysis, using the CoxPH function from the Survival package. Fecundity indices were analysed with ANOVAs fitted with the lm function in R.

## Acknowledgments

We thank Cathy Slack, Nazif Alic, Angela Douglas and Virginia Howick for comments which improved the manuscript. This study was funded by grants to MP from the Biotechnology and Biological Sciences Research Council (BBI011544/1) and the Royal Society (UF100158 & RG110303).

## Authors’ Contributions

AJD analysed data, performed experiments, designed the study and wrote the manuscript. XH performed experiments. E Blanc analyed data. E Bolukbasi contributed transgenic flies and performed experiments. YF performed experiments. MY performed experiments and designed the study. MDWP designed the study and wrote the paper.

## Data availability

RNAseq data have been deposited with NCBI. Custom R scripts have been uploaded to GitHub.

## Bibliography

1. Niccoli T, Partridge L. Ageing as a risk factor for disease. Curr Biol. 22, R741–752 (2012)

2. Piper MD, Partridge L, Raubenheimer D, Simpson SJ. Dietary Restriction and Aging: A Unifying Perspective. Cell Metab. 14 PAGE (2011).

3. McCay CM, Crowell MF, Marnard LA. The effect of retarded growth upon the length of life span and upon the ultimate body size. J Nutr. 10, 63–59 (1935).

4. Mair W, Piper MD, Partridge L. Calories Do Not Explain Extension of Life Span by Dietary Restriction in Drosophila. PLoS Bio. 3, e223 (2005).

5. Lee K, Simpson SJ, Clissold FJ, Brooks R, Ballard WJ, Taylor PW, et al. Lifespan and reproduction in Drosophila: New insights from nutritional geometry. Proc Natl Acad Sci. 7, 2498–2503 (2008).

6. Skorupa DA, Dervisefendic A, Zwiener J, Pletcher SD. Dietary composition specifies consumption, obesity, and lifespan in Drosophila melanogaster. Aging Cell 7, 478–490 (2008).

7. Grandison RC, Piper MD, Partridge L. Amino-acid imbalance explains extension of lifespan by dietary restriction in Drosophila. Nature 462, 1061–1064 (2009)

8. Solon-Biet SM, Aisling CM, Ballard WJ, Ruohonen K, Wu LE, Cogger VC, et al. The Ratio of Macronutrients, Not Caloric Intake, Dictates Cardiometabolic 631Health, Aging, and Longevity in Ad Libitum-Fed Mice. Cell Metab. 19, 418–430 (2014).

9. Solon-Biet SM, Mitchell SJ, Coogan SC, Cogger VC, Gokarn R, C M Aisling, et al. Dietary Protein to Carbohydrate Ratio and Caloric Restriction: Comparing Metabolic Outcomes in Mice. Cell Reports 11, 1529–1534 (2015).

10. Emran S, Yang M, He X, Zandveld J, Piper MD. Target of rapamycin signalling mediates the lifespan-extending effects of dietary restriction by essential amino acid alteration. Aging (Albany) 6, 390–8 (2014).

11. Simpson SJ, Couteur DG, Raubenheimer D. Putting the Balance Back in Diet. Cell 161, 1823 (2015).

12. Bar-Peled L, Sabatini D. Regulation of mTORC1 by amino acids. Trends Cell Biol. 24, 400–406 (2014)

13. Taylor RC, Dillin A. XBP-1 is a cell-nonautonoous regulator of stress resistance and longevity. Cell 153, 1435–1447 (2013).

14. Katewa S, Kapahi P. Role of TOR signaling in aging and related biological processes in Drosophila melanogaster. Exp Gerontol. 46, 382–390 (2011).

15. Vilchez D, Simic M, Dillin A. Proteostasis and aging of stem cells. Trends Cell Biol. 24, 161–171 (2014).

16. Bülow MH, Aebersold R, Pankratz MJ, Jünger MA. The Drosophila FoxA Ortholog Fork Head Regulates Growth and Gene Expression Downstream of Target of Rapamycin. PLoS ONE 5, e15171 (2010).

17. Robida-Stubbs S, Glover-Cutter K, Lamming D, Mizunuma M, Narasimhan S, Neumann-Haefelin E, et al. TOR signaling and rapamycin influence longevity by regulating SKN-1/Nrf and DAF-16/FoxO. Cell Metab. 15, 713–724 (2012).

18. Tiebe M, Lutz M, Garza A, Buechling T, Boutros M, Teleman AA. REPTOR and REPTOR-BP Regulate Organismal Metabolism and Transcription Downstream of TORC1. Dev Cell. 33, 272–84 (2015).

19. Alic N, Tullet JM, Niccoli T, Broughton S, Hoddinott MP, Slack C, et al. Cell-nonautonomous effects of dFOXO/DAF-16 in aging. Cell Rep. 6, 608–616 (2014).

20. Zhang P, Judy M, Lee S-J, Kenyon C. Direct and Indirect Gene Regulation by a Life-Extending FOXO Protein in C. elegans: Roles for GATA Factors and Lipid Gene Regulators. Cell Metab. 17, 85–100 (2013).

21. Potier D, Davie K, Hulselmans G, Sanchez M, Haagen L, Vân H-T, et al. Mapping Gene Regulatory Networks in Drosophila Eye Development by Large-Scale Transcriptome Perturbations and Motif Inference. Cell Reports 9, 2290–2303 (2014).

22. Dutta D, Dobson AJ, Houtz PL, Gläßer C, Revah J, Korzelius J, et al. Regional Cell-Specific Transcriptome Mapping Reveals Regulatory Complexity in the Adult Drosophila Midgut. Cell Reports 12, 346–358 (2015).

23. Langfelder P, Horvath S. WGCNA: an R package for weighted correlation network analysis. BMC Bioinformatics. 9, 559-PAGE (2008).

24. Mccormick M, Promislow D. Networks in the Biology of Aging: Powerful Tools for a Complex Process. Annu Rev Gerontol Geriat. 34, 243–266 (2014).

25. Langfelder P, Horvath S. Eigengene networks for studying the relationships between co-expression modules. Bmc Syst Biol. 1, 54 PAGE (2007).

26. Moore A, Jan L, Jan Y. Hamlet, a binary genetic switch between single-and multiple-dendrite neuron morphology. Science. 297, 1355–1358 (2002).

27. Moore AW, Roegiers F, Jan LY, Jan Y-NN. Conversion of neurons and glia to external-cell fates in the external sensory organs of Drosophila hamlet mutants by a cousin-cousin cell-type respecification. Genes Dev. 18, 623–628 (2004).

28. Cooper T. Transmitting the signal of excess nitrogen in Saccharomyces cerevisiae from the Tor proteins to the GATA factors: connecting the dots. FEMS Microbiol Rev. 26, 223–238 (2002).

29. Attardo GM, Higgs S, Klingler KA, Vanlandingham DL, Raikhel AS. RNA interference-mediated knockdown of a GATA factor reveals a link to anautogeny in the mosquito Aedes aegypti. Proc Natl Acad Sci. 100, 13374–13379 (2003).

30. Lossky M, Wensink P. Regulation of Drosophila yolk protein genes by an ovary-specific GATA factor. Mol Cell Biol. 15, 6943–6952 (1995).

31. Alic N, Giannakou ME, Papatheodorou I, Hoddinott MP, Andrews DT, Bolukbasi E, et al. Interplay of dFOXO and Two ETS-Family Transcription Factors Determines Lifespan in Drosophila melanogaster. Plos Genet. 10, e1004619 (2014).

32. Giannakou ME, Goss M, Jünger MA, Hafen E, Leevers SJ, Partridge L. Long-Lived Drosophila with Overexpressed dFOXO in Adult Fat Body. Science 305, 361 PAGE (2004).

33. Poirier L, Shane A, Zheng J, Seroude L. Characterization of the Drosophila gene-switch system in aging studies: a cautionary tale. Aging Cell. 7, 758–770 (2008).

34. Couteur DG, Samantha S-B, Cogger VC, Mitchell SJ, Senior A, de Cabo R, et al. The impact of low-protein high-carbohydrate diets on aging and lifespan. Cell Mol Life Sci 73, 1237–1252 (2016).

35. Sorrentino RP, Gajewski KM, Schulz RA. GATA factors in Drosophila heart and blood cell development. Semin Cell Dev Biol. 16, 107–116 (2005).

36. Sam S, Leise W, Hoshizaki D. The serpent gene is necessary for progression 706through the early stages of fat-body development. Mech Dev. 60, 197–205 (1996).

37. Murakami R, Okumura T, Uchiyama H. GATA factors as key regulatory molecules in the development of Drosophila endoderm. Dev Growth Differ. 47, 710581–589 (2005).

38. Buchon N, Osman D, David F, Fang H, Boquete J-P, Deplancke B, et al. Morphological and Molecular Characterization of Adult Midgut Compartmentalization in Drosophila. Cell Reports 3, 1725–1738 (2013).

39. Okumura T, Takeda K, Kuchiki M, Akaishi M. GATAe regulates intestinal stem 715cell maintenance and differentiation in Drosophila adult midgut. Dev Biol. 410, 71624–35 (2016).

40. Hansen IA, Attardo GM, Park J-H, Peng Q, Raikhel AS. Target of rapamycin-mediated amino acid signaling in mosquito anautogeny. Proc Natl Acad Sci. 101, 10626–10631 (2004).

41. Hansen IA, Attardo GM, Roy SG, Raikhel AS. Target of Rapamycin-dependent Activation of S6 Kinase Is a Central Step in the Transduction of Nutritional Signals during Egg Development in a Mosquito. J Biol Chem. 280, 20565–20572 (2005).

42. Park J-H, Attardo GM, Hansen IA, Raikhel AS. GATA Factor Translation Is the Final Downstream Step in the Amino Acid/Target-of-Rapamycin-mediated Vitellogenin Gene Expression in the Anautogenous Mosquito Aedes aegypti. J. Biol. Chem. 281, 11167–11176 (2006).

43. Martín D, Piulachs M, Raikhel A. A novel GATA factor transcriptionally represses yolk protein precursor genes in the mosquito Aedes aegypti via interaction with the CtBP corepressor. Mol Cell Biol. 21, 164–174 (2001).

44. Williams GC. Pleiotropy, natural selection, and the evolution of senescence. Evolution 11, 398–411 (1957).

45. Rera M, Azizi M, Walker D. Organ-specific mediation of lifespan extension: More than a gut feeling? Ageing Res Rev 12, 436–444 (2013).

46. Budovskaya YV, Wu K, Southworth LK, Jiang M, Tedesco P, Johnson TE, et al. An elt-3/elt-5/elt-6 GATA Transcription Circuit Guides Aging in C. elegans. Cell., 134, 1–13 (2007).

47. Murphy CT, A M Steven, Bargmann CI, Fraser A, Kamath RS, Ahringer J, et al. Genes that act downstream of DAF-16 to influence the lifespan of Caenorhabditis elegans. Nature. 424, 277–284 (2003).

48. Alic N, Andrews DT, Giannakou ME, Papatheodorou I, Slack C, Hoddinott MP, et al. Genome wide dFOXO targets and topology of the transcriptomic response to stress and insulin signalling. Mol Syst Biol. 7, 502 PAGE (2011).

49. Teleman AA. miR-200 De-FOGs Insulin Signaling. Cell Metab. 11, 8–9 (2010).

50. Slack C, Giannakou ME, Foley A, Goss M, Partridge L. dFOXO-independent effects of reduced insulin-like signaling in Drosophila. Aging Cell 10, 735–748 (2011).

51. Min K, Yamamoto R, Buch S, Pankratz M, Tatar M. Drosophila lifespan control by dietary restriction independent of insulin-like signaling. Aging Cell. 7, 199–206 (2008).

52. Kim J, Neufeld T. Dietary sugar promotes systemic TOR activation in Drosophila through AKH-dependent selective secretion of Dilp3. Nat Comms 6, 6846 PAGE (2015).

53. Buchon N, Osman D. All for one and one for all: Regionalization of the Drosophila intestine. Insect Biochem Mol Biol. 67:2–8 (2015).

54. Ren C, Webster P, Finkel SE, Tower J. Increased internal and external bacterial load during Drosophila aging without life-span trade-off. JOURNAL 6, 144–152 (2007).

55. Biteau B, Karpac J, Supoyo S, Matthew D, Lehmann R, Jasper H. Lifespan Extension by Preserving Proliferative Homeostasis in Drosophila. PLoS Genet. 6, e1001159 (2010).

56. Rera M, Bahadorani S, Cho J, Koehler C, Ulgherait M. Modulation of longevity and tissue homeostasis by the Drosophila PGC-1 homolog. Cell Metab. 14, 623–634 (2011).

57. Rera M, Clark R, Walker D. Intestinal barrier dysfunction links metabolic and inflammatory markers of aging to death in Drosophila. Proc Natl Acad Sci. 109, 21528–21533 (2012).

58. Ayyaz A, Li H, Jasper H. Haemocytes control stem cell activity in the Drosophila intestine. Nat Cell Biol. 17, 736–748 (2015).

59. Carlson K, Gardner K, Pashaj A, Carlson D. Genome-wide gene expression in relation to Age in large laboratory cohorts of Drosophila melanogaster. Genet Res Int, 835624 (2015)

60. Petkau K, Parsons B, Duggal A, Foley E. A deregulated intestinal cell cycle program disrupts tissue homeostasis without affecting longevity in Drosophila. J Biol Chem. 289, 28719–28729 (2014).

61. Wong A, Dobson AJ, Douglas AE. Gut microbiota dictates the metabolic response of Drosophila to diet. J Exp Biol. 217, 1894–1901 (2014).

62. Regan JC, Khericha M, Dobson AJ, Bolukbasi E, Rattanavirotkul N, Partridge L. Sex difference in pathology of the ageing gut mediates the greater response of female lifespan to dietary restriction. Elife. 5, e10956 (2016).

63. Hudry B, Khadayate S, Miguel-Aliaga I. The sexual identity of adult intestinal stem cells controls organ size and plasticity. Nature 530, 344–348 (2016).

64. Clark RI, Salazar A, Yamada R, Sorel F-G, Morselli M, Alcaraz J, et al. Distinct Shifts in Microbiota Composition during Drosophila Aging Impair Intestinal Function and Drive Mortality. Cell Rep. 12, 1656–1667 (2015).

65. Block DH, Kwame T-B, Kang H, Carlisle JA, Hanganu A, Lai T, et al. The Developmental Intestinal Regulator ELT-2 Controls p38-Dependent Immune Responses in Adult C. elegans. PLoS Genet. 11, e1005265 (2015).

66. Aronson BE, Stapleton KA, Krasinski SD. Role of GATA factors in development, differentiation, and homeostasis of the small intestinal epithelium. Am J Physiol Gastrointest Liver Physiol. 306, 474–490 (2014).

67. O’Brien LE, Soliman SS, Li X, Bilder D. Altered Modes of Stem Cell Division Drive Adaptive Intestinal Growth. Cell 147, 603–614 (2011).

68. Libert S, Chao Y, Chu X, Pletcher SD. Trade-offs between longevity and pathogen resistance in Drosophila melanogaster are mediated by NFκB signaling. Aging Cell 5, 533–43 (2006).

69. Bass T, Grandison RC. Wong R, Martinez P, Partridge L, Piper MDW. Optimization of dietary restriction protocols in Drosophila. J Gerontol A Biol Sci Med Sci 62, 1071–1081 (2007).

70. Metaxakis A, Partridge L. Dietary Restriction Extends Lifespan in Wild-Derived Populations of Drosophila melanogaster. PLoS ONE 8, e74681 (2013).

71. Kim D, Pertea G, Trapnell C, Pimentel H, Kelley R, Salzberg SL. TopHat2: accurate alignment of transcriptomes in the presence of insertions, deletions and gene fusions. Genome Biol 14, R36 PAGE (2013).

72. Anders S, J M Davis, Chen Y, Okoniewski M, Smyth GK, Huber W, et al. Count-based differential expression analysis of RNA sequencing data using R and Bioconductor. Nat Protoc. 8, 1765–1786 (2013).

73. Love MI, Huber W, Anders S. Moderated estimation of fold change and dispersion for RNA-seq data with DESeq2. JOURNAL 15 550 PAGE (2014).

74. Lex A, Gehlenborg N, Strobelt H. UpSet: visualization of intersecting sets. IEEE Trans Vis Comput Graph 20, 1983–1992 (2014).

75. Herrmann C, de Sande B, Potier D, Aerts S. i-cisTarget: an integrative genomics method for the prediction of regulatory features and cis-regulatory modules. JOURNAL 40, e114–e114 THIS ISN’T RIGHT (2012).

